# Intranasal ChAdOx1 nCoV-19/AZD1222 vaccination reduces shedding of SARS-CoV-2 D614G in rhesus macaques

**DOI:** 10.1101/2021.01.09.426058

**Authors:** Neeltje van Doremalen, Jyothi N. Purushotham, Jonathan E. Schulz, Myndi G. Holbrook, Trenton Bushmaker, Aaron Carmody, Julia R. Port, Claude K. Yinda, Atsushi Okumura, Greg Saturday, Fatima Amanat, Florian Krammer, Patrick W. Hanley, Brian J. Smith, Jamie Lovaglio, Sarah L. Anzick, Kent Barbian, Craig Martens, Sarah Gilbert, Teresa Lambe, Vincent J. Munster

## Abstract

Intramuscular vaccination with ChAdOx1 nCoV-19/AZD1222 protected rhesus macaques against pneumonia but did not reduce shedding of SARS-CoV-2. Here we investigate whether intranasally administered ChAdOx1 nCoV-19 reduces shedding, using a SARS-CoV-2 virus with the D614G mutation in the spike protein. Viral load in swabs obtained from intranasally vaccinated hamsters was significantly decreased compared to controls and no viral RNA or infectious virus was found in lung tissue, both in a direct challenge and a transmission model. Intranasal vaccination of rhesus macaques resulted in reduced shedding and a reduction in viral load in bronchoalveolar lavage and lower respiratory tract tissue. In conclusion, intranasal vaccination reduced shedding in two different SARS-CoV-2 animal models, justifying further investigation as a potential vaccination route for COVID-19 vaccines.

---

The severe acute respiratory syndrome coronavirus 2 (SARS-CoV-2) pandemic initiated the rapid development of vaccines based on a wide variety of platforms. Just 11 months later after the release of the first genome sequence, 13 vaccines are in phase III clinical trials and results of phase 3 clinical trial data for three different vaccines have been released^1–3^. These data suggest that vaccines based on the spike (S) protein of SARS-CoV-2, which generate a neutralizing antibody response, can reach an efficacy of up to 95%. Furthermore, several vaccines developed by Astrazeneca/Oxford, Bharat Biotech, CanSinoBIO, the Gamaleya Research Institute, Moderna/VRC, Pfizer/BioNTech, Sinopharm, Sinovac, and the Vector Institute have now been approved, fully or for emergency use.

In humans, most SARS-CoV-2 infections will present as asymptomatic or mild upper respiratory tract infection but are still accompanied by shedding of virus^4^. Depending on the study, shedding in asymptomatic infections was of shorter duration, but often to similar viral loads initially^4^. Asymptomatic as well as pre-symptomatic shedding has been associated with SARS-CoV-2 transmission^5–7^.

In preclinical non-human primate (NHP) challenge experiments, several vaccines were successful at preventing disease and reducing or preventing virus replication in the lower respiratory tract. However, subgenomic and genomic viral RNA was detected in nasal samples of all NHP experiments, dependent on vaccine dose^8–13^. Subgenomic viral RNA is indicative of replicating virus in the upper respiratory tract. It is currently unclear whether the detection of shedding in NHPs translate directly to humans.

It is possible that vaccination will result in attenuation or prevention of disease, but infection of the upper respiratory tract will occur even after vaccination possibly resulting in transmission. Currently, the majority of COVID-19 vaccines in development utilize an intramuscular (IM) injection, which predominantly produces a systemic IgG response and a poor mucosal response^14^. For a vaccine to elicit mucosal immunity, antigens will need to be encountered locally at the initial site of replication: the upper respiratory tract (URT).

Here, we evaluate the potential of using COVID-19 vaccine candidate ChAdOx1 nCoV-19 as an intranasal (IN) vaccine in the hamster and rhesus macaque models.

## Results

To evaluate the efficacy of an IN vaccination with ChAdOx1 nCoV-19, three groups of 10 Syrian hamsters^15^ were vaccinated with a single dose; group 1 received ChAdOx1 nCoV-19 via the IN route, group 2 received the same dose of vaccine via the IM route, and group 3 received control vaccine ChAdOx1 GFP via the IM route. Binding antibodies against SARS-CoV-2 S protein in peripheral blood were measured at −1 days post infection (DPI). Vaccination via either route resulted in high IgG titers (25,600-204,800) with no significant difference between vaccination routes (Figure 1A). Likewise, high neutralizing antibodies titers were detectable at −1 DPI. Intriguingly, neutralizing antibody titers were significantly higher in animals that received an IN vaccination (Figure 1B). For IN inoculation of Syrian hamsters 28 days post vaccination, we used isolate SARS-CoV-2/human/USA/RML-7/2020 which contains the D614G mutation in the S protein. Animals who received ChAdOx1 GFP started losing weight at 3 DPI and did not regain weight until 8 DPI. None of the vaccinated animals lost weight throughout the course of the experiment (Figure 1C). Six animals per group were swabbed daily up to 7 DPI. Viral RNA was detected in swabs from all animals. A significantly reduced amount of viral RNA was detected in nasal swabs from IN-vaccinated animals compared to control animals on 1-3 and 6-7 DPI. However, a significant reduction of viral RNA detected in oropharyngeal swabs from IM-vaccinated animals compared to control animals was only detected at 7 DPI (Mixed-effect analysis, p-value <0.05). When the area under the curve (AUC) was calculated as a measurement of total amount of viral RNA shed, IN-vaccinated animals shed significantly less than control animals (Kruskall-Wallis test, p=0.0074). Although viral RNA is an important measurement, the most crucial measurement in swabs is infectious virus. We found a significant difference between infectious virus detected in oropharyngeal swabs of IN-vaccinated animals compared to controls daily (Mixed-effect analysis, p-value<0.05). Likewise, the amount of infectious virus shed over the course of the experiment was significantly lower in IN-vaccinated animals than controls (Figure 1D, Kruskall-Wallis test, p=0.002). In contrast, we did not find a significant difference in AUC for viral RNA and infectious virus when comparing control and IM-vaccinated animals (Figure 1E). At five DPI, four animals in each group were euthanized. Viral load and infectious virus titer were high in lung tissue of control animals. We were unable to detect viral RNA or infectious virus in lung tissue from IN-vaccinated animals. Two animals in the IM group were weakly positive for genomic RNA, but not for subgenomic RNA or infectious virus (Figure 1F).

**Figure 1.**
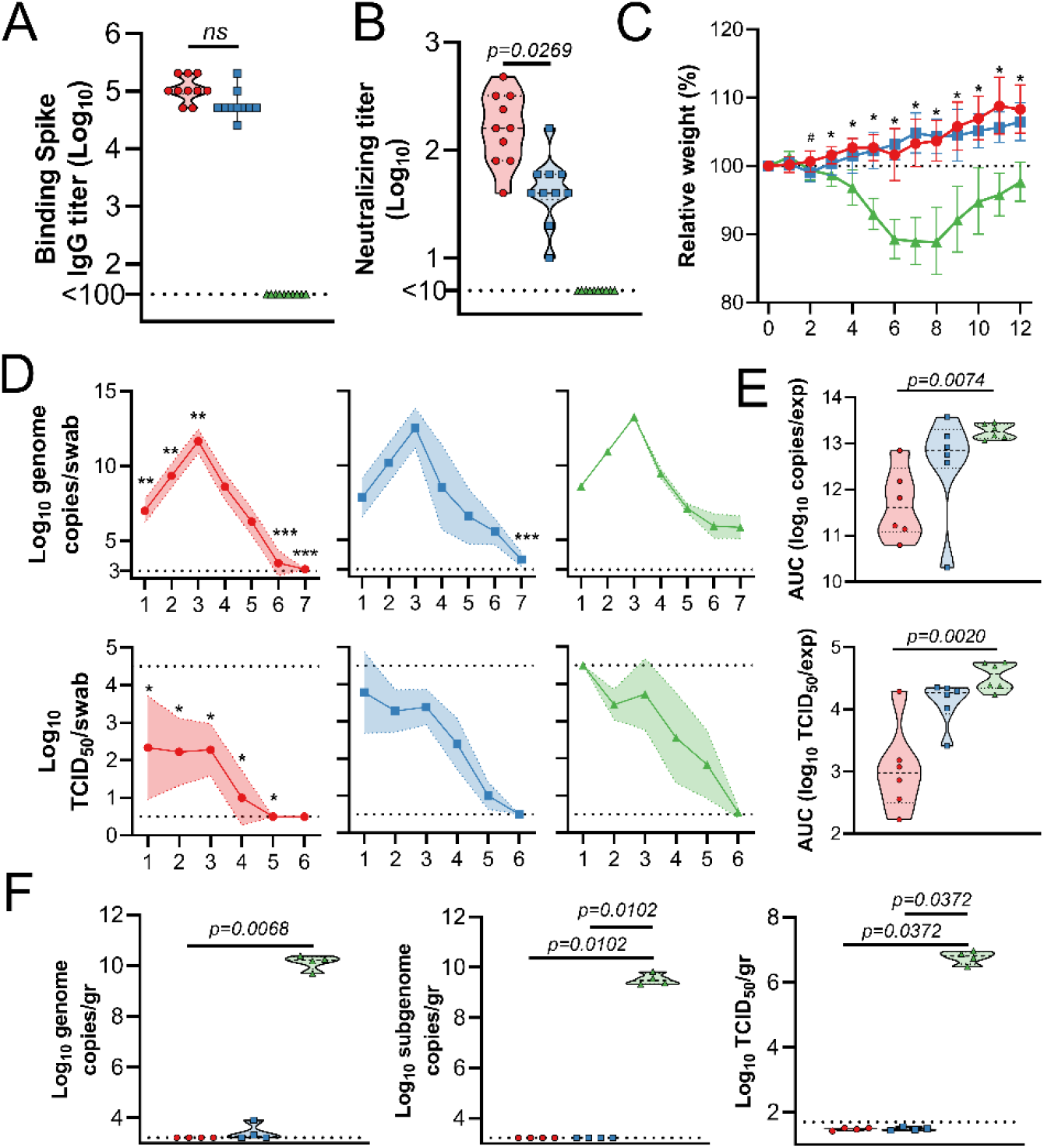
SARS-CoV-2 challenge of Syrian hamsters vaccinated with ChAdOx1 nCoV-19. Hamsters were vaccinated via the IN route (red), IM route (blue) or with control vaccine ChAdOx1 GFP via the IM route (green). A. Binding antibody titers against SARS-CoV-2 S protein. B. Virus neutralizing antibody titers. C. Relative weight upon challenge with SARS-CoV-2. A-B. Shown is geometric mean and 95% confidence interval. # = p-value <0.05 between IN and control group; * = p-value <0.05 between vaccinated groups and control group. D. Viral load and viral titer in oropharyngeal swabs. Shown is geometric mean (symbols) and 95% confidence interval (shade). E. Area under the curve analysis of viral load and titer shedding in oropharyngeal swabs. F. Viral load and titer in lung tissue, isolated at 5 DPI. E-F. Dashed line = median; dotted line = quartiles. Statistical analyses done using mixed-effect analyses (C), two-way ANOVA (D), or Kruskal-Wallis test (E-F). * = p value <0.05; ** = p-value < 0.01; *** = p-value < 0.001.

Lung tissue obtained at 5 DPI was then evaluated for pathology. Lesions were found in the lungs of control animals throughout (40-70% of tissue). Interstitial pneumonia was present in all animals, as well as edema, type II pneumocyte hyperplasia, and perivascular leukocyte infiltration, similar to what has been observed previously^15^. In stark contrast, no lesions or pathology were observed in lung tissue of vaccinated animals (Figure 2 and Table S1). SARS-CoV-2 N antigen in lung tissue was only found in control animals (20-70% of lung tissue was immunoreactive), but not for vaccinated animals (Figure 2 and Table S1).

**Figure 2.**
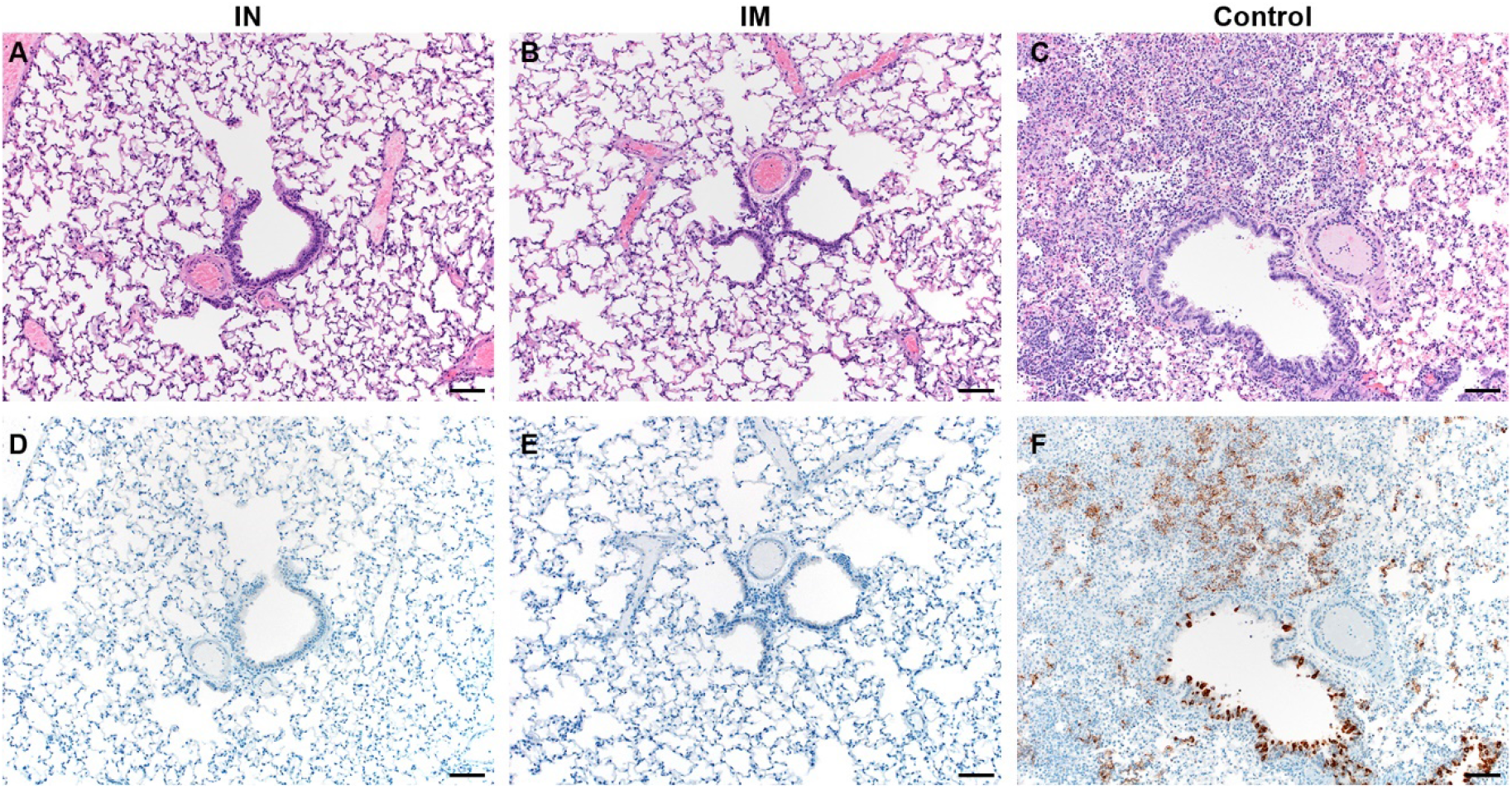
Pulmonary effects of direct intranasal challenge with SARS-CoV-2 in Syrian hamsters. A-C. H&E; D-F. IHC. A/B. No pathology. C. Moderate to marked interstitial pneumonia. D/E. No immunoreactivity. F. Numerous immunoreactive bronchiolar epithelial cells and Type I&II pneumocytes. Bar = 50µm.

Since direct IN inoculation of Syrian hamsters is an artificial route of virus challenge, and Syrian hamsters transmit SARS-CoV-2 readily^16^, we repeated the above experiment within a direct contact horizontal transmission setting. Briefly, unvaccinated hamsters were IN challenged with SARS-CoV-2 (donor animals). After 24 hours, vaccinated animals were introduced into the cage, then four hours later, donor animals were removed (Figure 3A). As in the previous experiment, vaccination of hamsters with ChAdOx1 nCoV-19 resulted in high binding and neutralizing antibodies. Neutralizing antibodies were significantly higher in IN-vaccinated animals (Figure 3B-C). Control animals started losing weight at 4 days post exposure (DPE) and started recovering weight at 8 DPE. None of the vaccinated animals lost weight throughout the experiment, and a significant difference in weight was observed starting at 4 and 5 DPE for IN and IM-vaccinated animals compared to controls, respectively (Figure 3D). Ten animals per group were swabbed daily. Shedding of viral RNA, but not infectious virus, in controls was lower than in the previous experiment (multiple unpaired two-tailed Student’s t-test, 2-4 and 7 DPI, p-value <0.0001). A significantly reduced amount of shedding, both for viral RNA and infectious virus, was again detected in IN-vaccinated animals. However, as in the previous experiment, limited significant differences in shedding were detected in IM-vaccinated animals compared to controls (Figure 3E, mixed effects, p<0.05). The total amount of shedding, illustrated as AUC, was significantly different for IN-vaccinated animals compared to controls in both viral RNA and infectious virus (Kruskal-Wallis, p-value <0.0001), but not for IM-vaccinated animals (Figure 3F). Four animals per group were euthanized at 5 DPE and lung tissue was harvested. Again, no viral RNA or infectious virus was detected in lung tissue obtained from IN-vaccinated animals. However, viral RNA could be detected in lung tissue from three (gRNA) and two (sgRNA) IM-vaccinated animals, whereas infectious virus was detected in lung tissue of one IM animal (Figure 3G).

Virus obtained from oropharyngeal swabs was sequenced at 2 and 5 DPE. Sequences obtained at 2 DPE from four different animals contained SNPs in the S protein. Two SNPs encoded a non-synonymous mutation; Asp839Glu and Lys1255Gln. Three swabs were obtained from IN-vaccinated animals, one swab was obtained from an IM-vaccinated animal (Table 1).

**Table 1.** SNPs in SARS-CoV-2 sequences obtained from hamster swabs.

| Nucleotide change | Amino acid change | Group | Number of reads Mutation/total (%) | Day |
| --- | --- | --- | --- | --- |
| A23911T | Ala783Ala | IN | 243/260 (93.5) | 2 DPE |
| T24079G | Asp839Glu | IN | 250/391 (63.9) | 2 DPE |
| A24253C | Pro897Pro | IN | 425/641 (66.3) | 2 DPE |
| A25325C | Lys1255Gln | IM | 273/768 (35.5) | 2 DPE |

**Figure 3.**
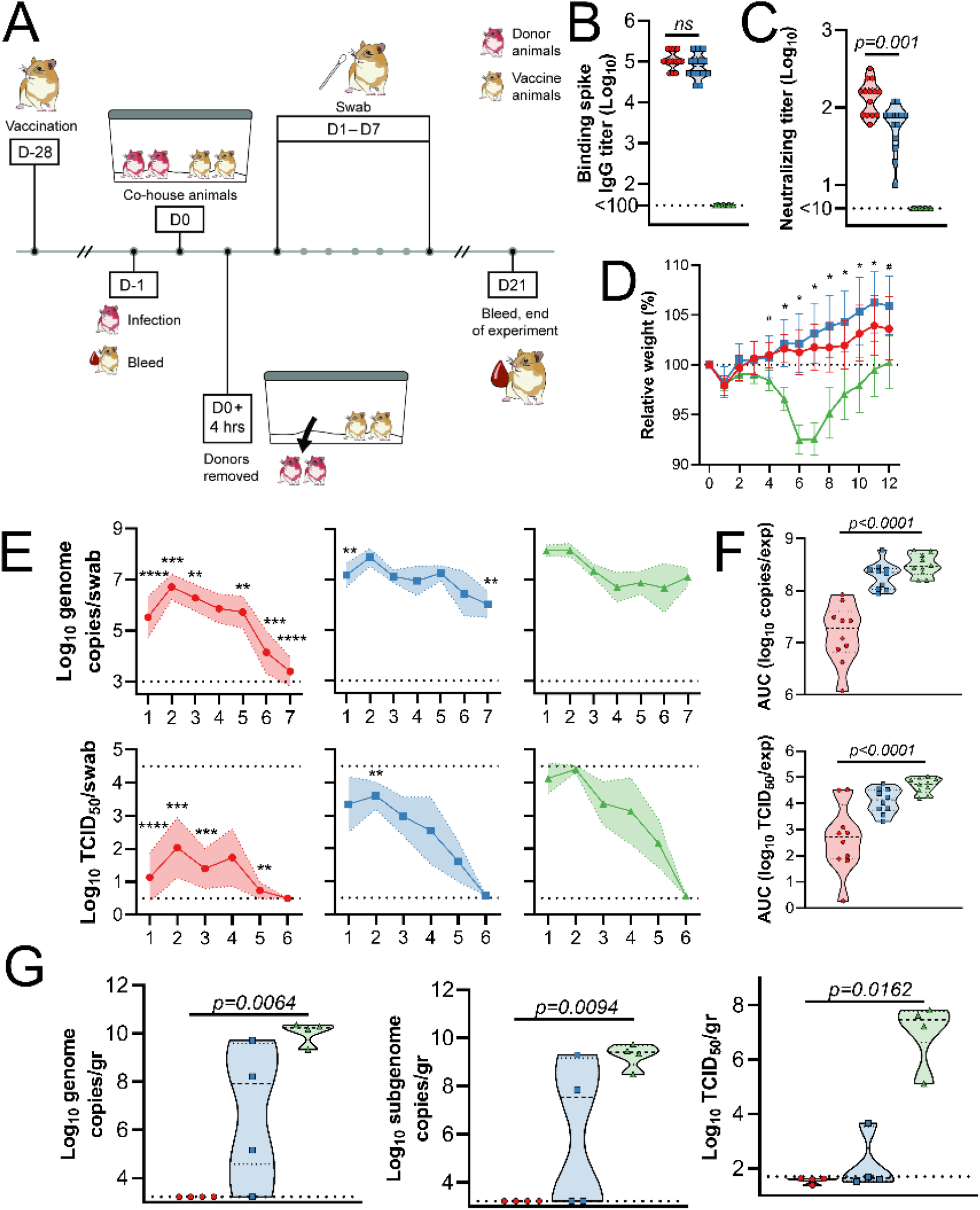
SARS-CoV-2 transmission to Syrian hamsters vaccinated with ChAdOx1 nCoV-19. Hamsters were vaccinated via the IN route (red), IM route (blue) or with control vaccine ChAdOx1 GFP via the IM route (green). A. Experimental schedule. Hamsters received a single vaccination 28 days prior to exposure. Donor animals were challenged at −1 DPE, and hamsters were co-housed for 4 hours. one day later. B. Binding antibody titers against SARS-CoV-2 S protein. C. Virus neutralizing antibody titers. D. Relative weight upon challenge with SARS-CoV-2. Shown is geometric mean and 95% confidence interval. # = p-value <0.05 between IN and control group; * = p-value <0.05 between vaccinated groups and control group. E. Viral load and viral titer in oropharyngeal swabs. Shown is geometric mean (symbols) and 95% confidence interval (shade). F. Area under the curve analysis of viral load and titer shedding in oropharyngeal swabs. G. Viral load and titer in lung tissue, isolated at 5 DPE. F-G. Dashed line = median; dotted line = quartiles. Statistical analyses done using mixed-effect analyses (C), two-way ANOVA (D), or Kruskal-Wallis test (E-F). * = p value <0.05; ** = p-value < 0.01; *** = p-value < 0.001.

Lung tissue of control animals obtained at 5 DPE had the same appearance as those obtained in the previous experiment. Lesions were observed in 40-50% of tissue, and interstitial pneumonia, edema, type II pneumocyte hyperplasia, and perivascular leukocyte infiltration were observed in all animals. As previously, no lesions or pathology were observed in lung tissue of IN-vaccinated animals. However, lesions were observed in the IM-vaccinated animals (5-20%, 3 out of 4 animals), accompanied with mild interstitial pneumonia (3 out of 4 animals), type II pneumocyte hyperplasia (2 out of 4 animals), and perivascular leukocyte infiltration (1 out of 4 animals). Edema was not observed (Figure 4 and Table S2). SARS-CoV-2 N antigen in lung tissue was found to be present in control animals (30-60% of lung tissue was immunoreactive) and to a lesser extent in IM-vaccinated animals (5% of lung tissue, 3 out of 4 animals), but not for IN-vaccinated animals (Figure 4 and Table S2).

**Figure 4.**
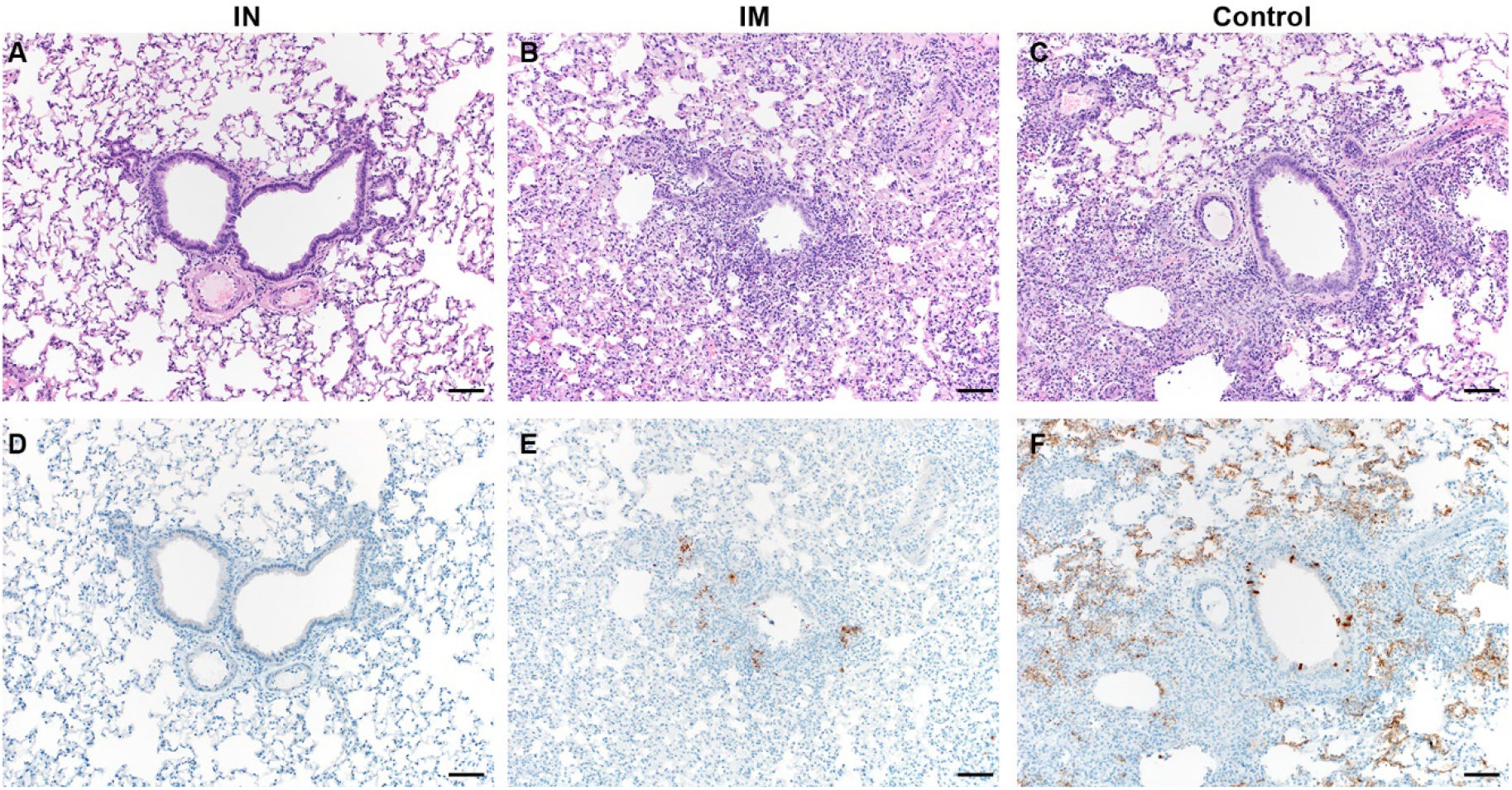
Pulmonary effects of transmission of SARS-CoV-2 in Syrian hamsters. A-C. H&E; D-F. IHC. A. No pathology. B. Mild interstitial pneumonia. C. Moderate to marked interstitial pneumonia. D. No immunoreactivity. E. Scattered immunoreactive bronchiolar epithelial cells and Type I&II pneumocytes. F. Numerous immunoreactive bronchiolar epithelial cells and Type I&II pneumocytes. Bar = 50µm

The results obtained in hamster studies prompted us to investigate an IN vaccination in rhesus macaques^17^. Four non-human primates were vaccinated with a prime-boost regimen of ChAdOx1 nCoV-19 using the same dose as previously described^8^, utilizing an IN mucosal atomization device, which produced a spray of aerosols that were deposited in the nasal cavity. Four control animals were vaccinated with ChAdOx1 GFP. Blood, nasosorption swabs and bronchoalveolar lavage (BAL) samples were collected throughout the experiment. As expected, a higher fraction of IgA to total Ig was detected in nasosorption samples compared to BAL and serum samples (Figure S1). S and RBD-specific IgG antibodies were detected in serum and nasosorption samples after prime vaccination, but not in BAL, at seven days post prime vaccination (−49 DPI). Higher IgG titers were found in all samples obtained after a second vaccination at −28 DPI (Figure 5A-C). S and RBD-specific IgA antibodies were detected in serum upon prime vaccination but did not increase upon boost vaccination (Figure 5D). In contrast, SARS-CoV-2 specific IgA antibodies were only weakly detected in nasosorption samples upon prime vaccination but further increased upon boost vaccination (Figure 5E). No SARS-CoV-2 specific IgA antibodies were detected in BAL at −49 DPI but were detected seven days post boost vaccination (−21 DPI, Figure 5F). Circulating neutralizing antibodies were readily detected in vaccinated animals, to levels similar to convalescent serum obtained from human survivors (varying from asymptomatic to severe) and from NHPs which received a prime or prime-boost IM vaccinated with ChAdOx1 nCoV-19^8^ (Kruskal-Wallis test, Figure 5G). Furthermore, multiple antigen-specific antibody Fc effector functions were detected; circulating antibodies in vaccinated animals promoted phagocytosis, complement deposition and NK cell activation (Figure 5H). Finally, levels of binding antibodies at 0 DPI against wildtype RBD and N501Y RBD were compared. N501Y is found in the RBD of two new variants of SARS-CoV-2: VOC2020-12/01 (B.1.1.7) and 501Y.V2 (B.1.351). No differences were found in the level of binding antibodies between wildtype and 501Y mutant RBD (Figure 5I).

**Figure 5.**
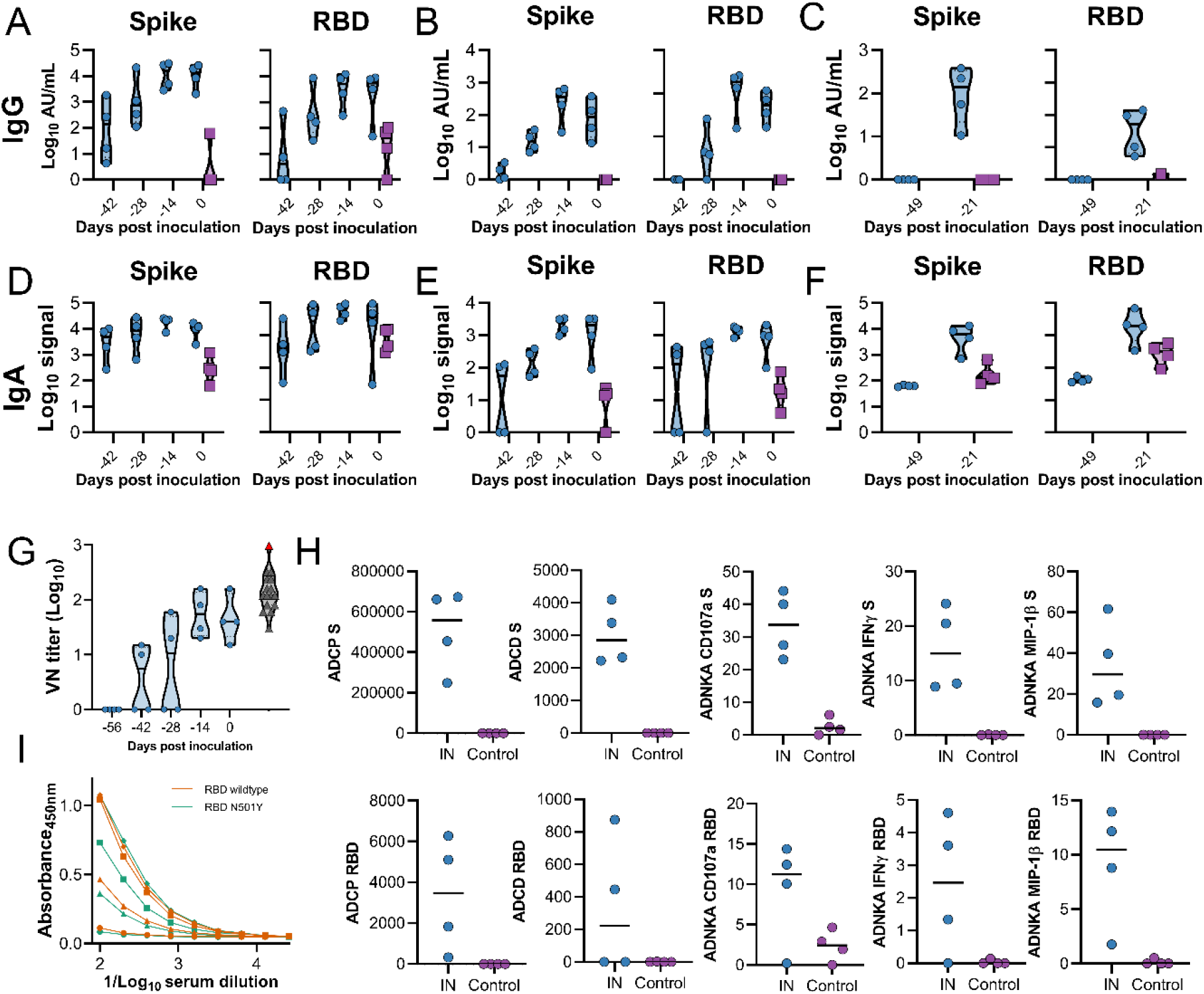
Humoral response to IN vaccination with ChAdOx1 nCoV-19 in rhesus macaques. Truncated violin plot of SARS-CoV-2-specific IgG antibodies measured in serum (A), nasosorption samples (B), and BAL (C). Truncated violin plot of SARS-CoV-2-specific IgA antibodies measured in serum (D), nasosorption samples (E), and BAL (F). G. Truncated violin plot of neutralizing antibodies in serum. H. Effector functions of antibodies in serum. Black line = median; dotted line = quartiles; blue = vaccinated animals; purple = control animals (only 0 DPI values shown); black = human convalescent sera; red = NIBSC serum control 20/130. I. Anti-RBD-specific IgG titers at 0 DPI. Orange = RBD wildtype; aqua = RBD N501Y.

Animals were challenged via the intratracheal and IN route using 10^6^ TCID_50_ of SARS-CoV-2 (SARS-CoV-2/human/USA/RML-7/2020). Nasal swabs were investigated for the presence of genomic RNA, subgenomic RNA and infectious virus. In control animals, both types of viral RNA were readily detected in nasal swabs. Genomic RNA was detected in all 4 animals (11 out of 16 swabs total), whereas subgenomic RNA was detected in 3 out of 4 animals (4 out of 16 swabs total). Infectious virus was detected in 3 out of 4 animals (5 out of 16 swabs total). Viral RNA was detected in nasal swabs obtained from vaccinated animals, but viral load was lower and fewer swabs were positive. Genomic RNA was detected in 3 out of 4 animals (5 out of 16 swabs total), whereas subgenomic RNA and infectious virus was only detected in 1 out of 4 animals (1 swab each) (Figure 6A). Total amount shed was depicted using AUC analysis. Although a downwards trend was observed in nasal swabs from vaccinated animals, this difference was not statistically significant (Mann-Whitney test, Figure 6B). Genomic and subgenomic RNA in BAL was detected in all four control animals (11 and 8 out of 12 samples, respectively). Infectious virus in BAL was detected in 2 out of 4 animals (3 out of 8 samples). Genomic RNA was detected in 4 out of 4 vaccinated animals, but only at early time points (5 out of 12 samples). Subgenomic RNA was only found in one animal and was very low (1 out of 12 samples). The differences in number of positive samples between vaccinated and control animals were significant using a Fisher’s exact test (genomic RNA p-value = 0.0272; subgenomic RNA p-value = 0.0094). No infectious virus could be detected in BAL samples from vaccinated animals (0 out of 12 samples) (Figure 6C). AUC analyses again showed a downwards trend in BAL from vaccinated animals, which was significant for sgRNA (Figure 6D, p=0.0286, Mann-Whitney test). Animals were euthanized at 7 DPI and viral RNA in nasal turbinates and lung tissue was analyzed. Viral load in lung was significantly lower for vaccinated animals than for control animals (Mann-Whitney test, p-value <0.0001 and 0.001 for genomic and subgenomic RNA, respectively), but no difference in viral load in nasal turbinates was detected (Figure 6E-F). No differences in hematology and radiographs were observed between groups.

**Figure 6.**
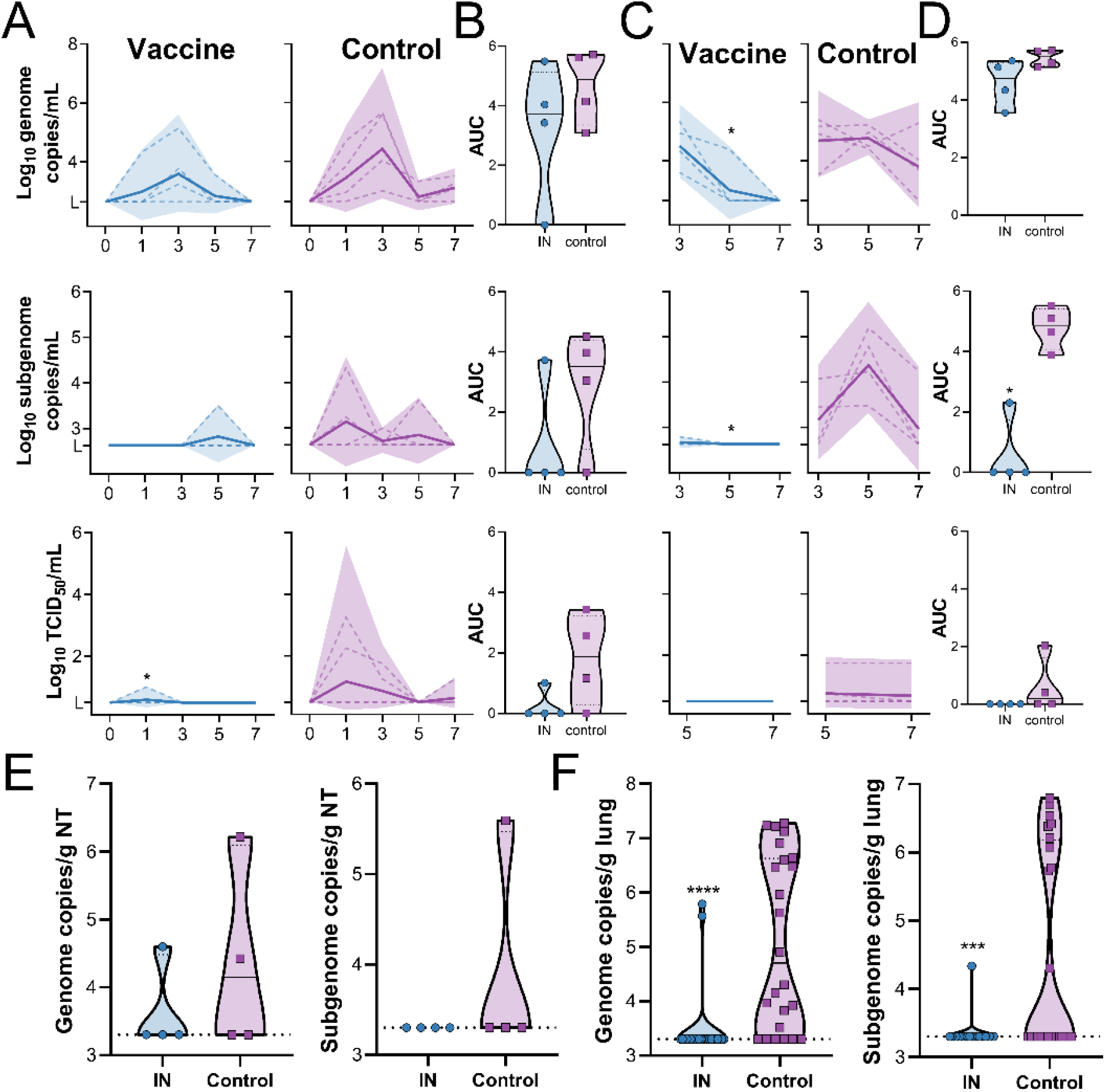
SARS-CoV-2 detection in samples obtained from rhesus macaques upon virus challenge. gRNA, sgRNA and infectious virus in nasal swabs (A) and BAL (C) was determined. Dotted line = individual animals; solid line = geometric mean; shaded area = 95% confidence interval. Area under the curve (AUC) was calculated as an indication of the total amount of virus shed in nasal swabs (B) and BAL (D) and displayed as a truncated violin plot. Solid line = median; dotted line = quartiles. * = p-value <0.05 as determined via two-tailed Mann-Whitney test. Amount of gRNA and sgRNA in nasal turbinate (E) and lung tissue (F). Blue = vaccinated animals; purple = control animals; solid line = median; dotted line = quartiles. *** = p-value <0.001; **** = p-value <0.0001, as determined via two-tailed Mann-Whitney test.

We subsequently sought to define the impact of the vaccine-specific humoral response on nasal shedding and viral load after challenge. Principal component (PC) analysis of the pre-challenge, multivariate antibody profile revealed the distinct segregation of vaccinated animals from controls, driven by local and systemic antibodies with diverse functions (Figures 7A and 7B). Intergroup variation, largely encapsulated by PC2, was primarily mediated by differences in virus-specific IgA or IgG antibody levels in BAL and nasosorption samples. Notably, minimal levels of nasosorption IgG and relatively low levels of nasosorption IgA were detected in the only animal exhibiting significant nasal shedding after challenge (NHP1). This animal also had low serum IgG and virus neutralizing titers. Meanwhile, levels of BAL IgA and IgG were lowest in NHP2 and very high in NHP4; genomic and subgenomic RNA levels in BAL and lung tissue were highest and lowest in these animals, respectively. PC analysis of post-challenge viral load (AUC) in nasal swabs, BAL, and lung tissue, again, yielded dramatic clustering according to vaccination status (Figure 7C). Variation between control animals seemed to reflect site-specific differences in virus replication, between the upper and lower respiratory tract (Figure 7D).

**Figure 7.**
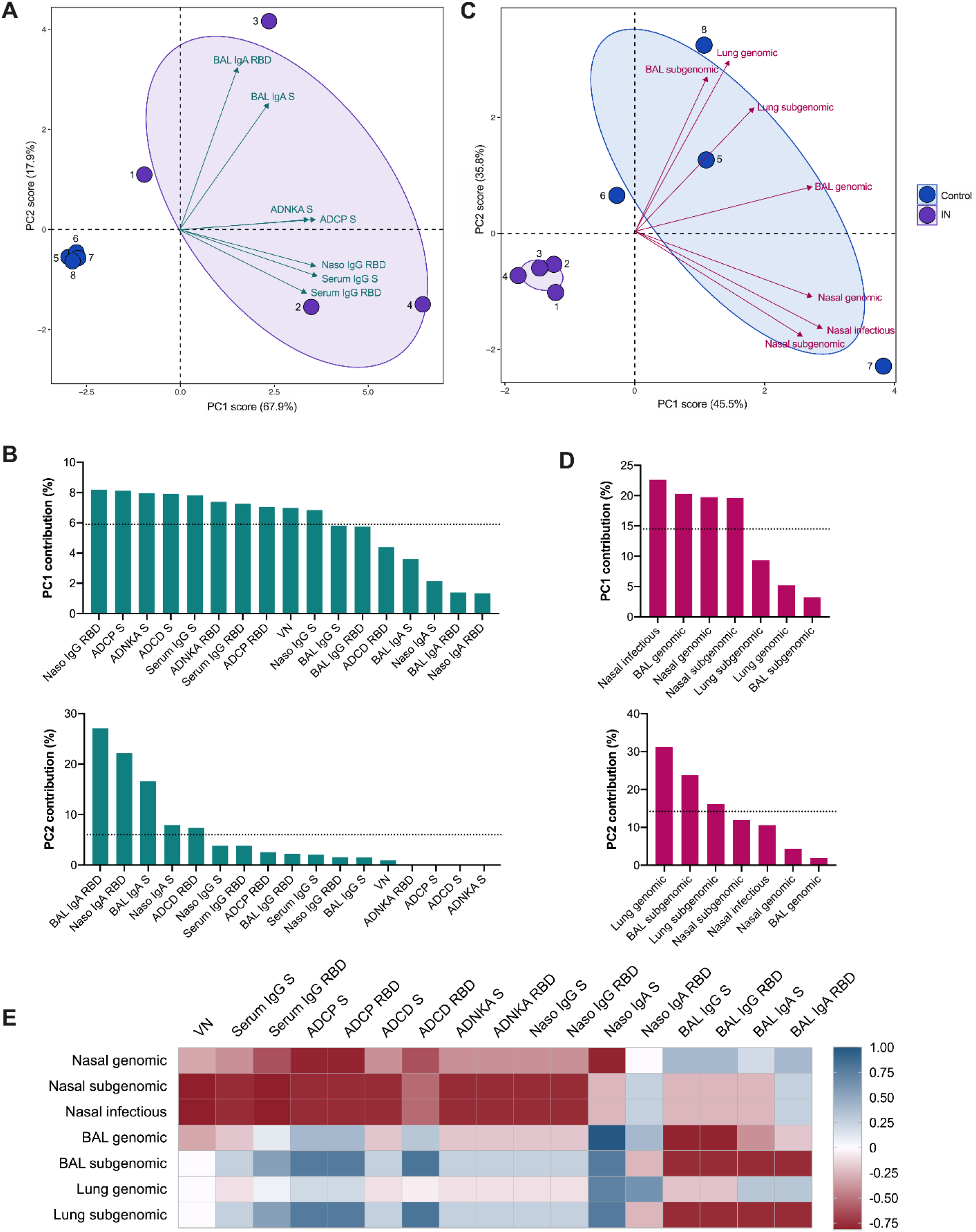
Influence of the vaccine-induced humoral response on viral RNA levels post challenge. Principal component analysis (PCA) plot of the multivariate antibody (A) and AUC virology (C) profile across all animals (numbered dots). Ellipses indicate group distribution as 95% confidence levels. Mapped arrow projections indicate the influence of individual variables on the PCs; the antibody plot depicts only the top seven contributors. The complete antibody (B) and virology (D) variable loading plots for PCs 1 and 2 with a dotted line to indicate average expected contribution. Heatmap visualization of the correlations between antibody measures and viral RNA (AUC) levels (E) for the IN-vaccinated animals; R values were generated using two-sided Spearman rank correlation tests.

To examine these relationships further, we generated a correlation matrix integrating the pre- and post-challenge data from the IN-vaccinated animals (Figure S2). The Spearman rank correlation coefficients computed for individual antibody-virology variable pairings were assessed; however, the low number of animals precluded observations of statistical significance (Figure 7E). Nevertheless, clear trends emerged. While higher levels of serum (neutralizing and Fc effector-function-inducing) antibodies and nasosorption antibodies correlated with reduced nasal shedding, viral RNA in the BAL and lung tissue exclusively displayed strong negative correlations with BAL IgG and IgA levels. Of note, subgenomic RNA levels generally appeared to correlate more strongly with antibody levels than genomic RNA levels across sampling sites.

## Discussion

Here we show that IN vaccination of hamsters and NHPs with ChAdOx1 nCoV-19 results in a robust mucosal and humoral immune response. In comparison to hamsters vaccinated via the IM route, a reduction in shedding is found in IN vaccinated animals, combined with full protection of the respiratory tract (no viral RNA). In NHPs, we observed a reduction in nasal shedding and viral load in BAL, as well as protection of the lower respiratory tract.

Since the release of the first full-length genome of SARS-CoV-2^18^, thousands of complete genomes have been released. Multiple clades have been identified, as well as mutations throughout the genome of SARS-CoV-2. The most prevalent of the novel mutations is likely D614G in the S protein, which is present in the majority of circulating SARS-CoV-2 viruses^19^. All vaccines in clinical trials are based on the initial SARS-CoV-2 sequences^18^, and mutations in the S protein may result in immune evasion^20^. Here, a heterologous challenge was implemented in all experiments; we utilized isolate SARS-CoV-2/human/USA/RML-7/2020, which was isolated from a nasopharyngeal swab in July 2020 and belongs to clade 20A. This virus has one coding change in the S protein compared to the vaccine antigen; D614G. Both hamster and NHP studies described here demonstrate clearly that the ChAdOx1 nCoV-19 vaccine protects against SARS-CoV-2 containing the D614G mutation. It is likely that this translates to other vaccine platforms as well. Recently, new variants of SARS-CoV-2, named VOC2020-12/01 (B.1.1.7) and 501Y.V2 (B.1.351) were detected^21^ and VOC2020-12/01 (B.1.1.7) was potentially linked to increased transmission^22^. Both strains contain the N501Y mutation in RBD. We detected no difference in the ability of antibodies elicited by ChAdOx1 nCoV-19 vaccination to bind to N501Y mutant RBD.

Our previous and others’ studies investigating efficacy of COVID-19 vaccines in NHPs showed complete or near complete protection of the lower respiratory tract, but nasal shedding was still observed^8–13^. In natural infection with respiratory pathogens, a systemic immune response, dominated by IgG, as well as a mucosal immune response, dominated by secretory IgA (sIgA), is induced^14,23^. Although abundant literature exists on systemic immune responses to natural SARS-CoV-2 infection, literature on mucosal immunity is currently limited. In mucosal fluids from COVID-19 patients, S and RBD-specific IgA, IgG, and IgM were readily detected^24–26^. It is hypothesized that sIgA mainly protects the upper respiratory tract, whereas systemic IgG protects the lower respiratory tract^14,27,28^.

Upon IN vaccination of rhesus macaques with ChAdOx1 nCoV-19, we were able to detect SARS-CoV-2 specific IgG and IgA in serum. More importantly, SARS-CoV-2 specific IgG and IgA was also detected in nasosorption samples and BAL. No nasosorption samples were collected in our previous study^8^, but BAL collected at 3 and 5 DPI did not contain high levels of SARS-CoV-2 specific antibodies. Thus, IN vaccination resulted in systemic immunity comparable to that induced in vaccinated animals who received an IM vaccination with ChAdOx1 nCoV-19, but also elicited SARS-CoV-2-specific mucosal immunity as demonstrated by IgA detection in nasosorption and BAL samples.

Mucosal vaccination resulted in a reduction in shedding. In NHPs, subgenomic and infectious virus shedding was only detected in one vaccinated animal. This animal exhibited low levels of IgG and IgA antibodies in nasosorption samples coupled with low VN and sera IgG titers, suggesting that a robust humoral response in the nasal mucosa and in circulation is necessary to efficiently control nasal shedding.

Vaccination of small animal models with an adeno-vectored vaccine against SARS-CoV-2 has been reported by others, including two studies which investigated IN vaccination^29–31^. Bricker *et al*. showed a reduction in nasal shedding, near complete protection of upper respiratory tract and partial lower respiratory tract protection in hamsters^29^, whereas Hassan *et al*. did not investigate nasal shedding but found near complete protection of upper and lower respiratory tract tissue in mice^30^. Tostanoski *et al*. investigated IM vaccination in hamsters and found near complete protection in lung tissue dependent on vaccine candidate^31^. This agrees with our findings; we find a reduction in nasal shedding in IN-vaccinated animals, but not IM-vaccinated animals. We also find full protection of the lower respiratory tract in IN-vaccinated animals. Since IN vaccination of mice^30^ and NHPs elicited SARS-CoV-2-specific IgA in BAL and nasosorption samples (NHP only), we hypothesize that the same occurred in hamsters and combined with the higher neutralizing titers resulted in a reduction in nasal shedding.

In our second hamster study we moved away from IN inoculation and investigated vaccine efficacy in a transmission model. Transmission of SARS-CoV-2 was efficient, resulting in 100% transmission to control sentinel animals after just 4 hours of exposure to infected animals. Again, IN vaccination resulted in a reduction in shedding in sentinel hamsters compared to control animals. Although protection of the lower respiratory tract was complete in IN-vaccinated animals, only partial control was seen in IM-vaccinated animals, in contrast to the direct challenge experiment. It is possible that the difference between IN and IM-vaccinated animals is caused by virus seeding of the lungs from the URT; higher viral nasal shedding in IM-vaccinated animals compared to IN-vaccinated animals is likely reflective of a relative increase in virus deposition in the lung from the URT in IM compared to IN-vaccinated animals. That does not explain however why such a discrepancy between vaccination groups was not observed in the direct challenge study. Another hypothesis would be a difference in the initial site of virus deposition. Direct contact transmission likely represents a wide variety of exposure routes for the sentinel animals, including fomites and aerosols. A previous study in our laboratory showed the deposition of fluorescently labeled virus in the lungs of hamsters upon IN inoculation^32^. However, whereas that study used an inoculation volume of 80 µl, in the current study an inoculation volume of 40 µl was utilized, and it is possible that virus deposition directly into the lungs via IN inoculation with 40 µl is limited, whereas in direct transmission virus particles might be inhaled directly into the lung. Indeed, we recently showed that infection via aerosols, but not via direct IN inoculation, resulted in a high virus load in lung tissue at 1 DPI^33^. It should be noted that infectious virus titers in the lungs of control animals in the transmission experiment compared to the direct challenge experiment were ∼5x higher, supporting this supposition. Mercado *et al*. previously showed the importance of different effector functions of antibodies in protection against SARS-CoV-2 in rhesus macaques^13^. We adapted their assays and show that upon IN vaccination with ChAdOx1 nCoV-19, a variety of antibody-dependent Fc effector functions are elicited, including monocyte cellular phagocytosis, complement deposition, and natural killer cell activation. Although the importance of neutralizing antibodies against SARS-CoV-2 has been convincingly demonstrated in rhesus macaques^34^, the importance of other effector functions remains unknown. ChAdOx1 nCoV-19 has been shown to induce anti-S neutralizing antibody titers, as well as antibody-dependent neutrophil/monocyte phagocytosis, complement activation and natural killer cell activation^35^. A selective delay or defect in IgG development has been linked to severe and fatal outcome in human patients^36^. A recent study in mice demonstrated that *in vitro* neutralization did not uniformly correlate with *in vivo* protection, and that binding to Fc receptors was of importance, suggesting that antibody effector functions play a pivotal role in protection against SARS-CoV-2^37^. Preliminary PC and correlation analyses suggested that while both vaccine-induced circulating antibodies—with neutralizing and non-neutralizing functionality—and upper respiratory antibodies play a role in reducing nasal shedding, virus replication in the airway and lung tissue is primarily controlled by antibodies localized to the lower respiratory tract. However, given that low animal numbers prevented us from establishing correlations of statistical significance, further studies will be required to more clearly define the relative impact of each component of the multifunctional humoral response on measures of protection.

The data presented supports the investigation of IN delivery of COVID-19 vaccines. With the roll-out of COVID-19 vaccines worldwide, it will be crucial to investigate whether the vaccines provide sterilizing immunity, or whether vaccinated people are still susceptible to infection of the URT and onward transmission of the virus. The data presented here demonstrates SARS-CoV-2-specific mucosal immunity is possible after IN vaccination, and results in a reduction in nasal shedding. It is now pertinent to investigate whether this finding translates to the clinic.

## Acknowledgments

We thank O. Abiona, B. Bailes, R. Cole, K. Corbett, K. Cordova, L. Crawford, H. Feldmann, B. S. Gallogly, Graham, L. Heaney, M. Jones, R. LaCasse, M. Marsh, K. Menk, A. Mora, Rose Perry, R. Rivera, L. Shupert, B. Smith, A. Weidow, and M. Woods for their assistance during this study.

## Author contributions

N.v.D., J.R.P., and V.M. designed the studies, N.v.D., J.N.P., J.E.S., M.G.H., T.B., A.C., J.R.R., C.K.Y, A.O., G.S., J.L., P.W., B.S., S.L.A., K.B., and V.J.M. performed the experiments, F.A. and F.K. designed and provided RBD protein, N.v.D, J.N.P, A.O., J.L., P.H., S.L.A., C.M., S.G., T.L., and V.J.M analyzed results, N.v.D and J.N.P wrote the manuscript, all co-authors reviewed the manuscript.;

## Data and materials availability

All data is available in the main text or the supplementary materials.

## Supplementary Materials

### Materials and Methods

#### Ethics statement

The Institutional Animal Care and Use Committee (IACUC) at Rocky Mountain Laboratories provided all animal study approvals, which were conducted in an Association for Assessment and Accreditation of Laboratory Animal Care (AAALAC)-accredited facility, following the basic principles and guidelines in the Guide for the Care and Use of Laboratory Animals 8^th^ edition, the Animal Welfare Act, United States Department of Agriculture and the United States Public Health Service Policy on Humane Care and Use of Laboratory Animals.

Animals were kept in climate-controlled rooms with a fixed light/dark cycle (12-hours/12-hours). Hamsters were co-housed in rodent cages, fed a commercial rodent chow with ad libitum water and monitored at least once daily. Rhesus macaques were housed in individual primate cages allowing social interactions, fed a commercial monkey chow, treats and fruit with ad libitum water and were monitored at least twice daily. Environmental enrichment for rhesus macaques consisted of a variety of human interaction, commercial toys, videos, and music. The Institutional Biosafety Committee (IBC) approved work with infectious SARS-CoV-2 virus strains under BSL3+ conditions. All sample inactivation was performed according to IBC approved standard operating procedures for removal of specimens from high containment.

#### Generation of ChAdOx1 nCoV-19 vaccine

ChAdOx1 nCoV-19 was designed as previously described^8^. Briefly, the S protein of SARS-CoV-2 (GenBank accession number YP_009724390.1) was codon optimized for expression in human cell lines and synthesized with the tissue plasminogen activator (tPA) leader sequence at the 5’ end by GeneArt Gene Synthesis (Thermo Fisher Scientific). The sequence, encoding SARS-CoV-2 amino acids 2-1273 and tPA leader, was cloned into a shuttle plasmid using InFusion cloning (Clontech). The shuttle plasmid encodes a modified human cytomegalovirus major immediate early promoter (IE CMV) with tetracycline operator (TetO) sites, poly adenylation signal from bovine growth hormone (BGH). ChAdOx1 nCoV-19 was prepared using Gateway® recombination technology (Thermo Fisher Scientific) between this shuttle plasmid and the ChAdOx1 destination DNA BAC vector^38^ resulting in the insertion of the SARS-CoV-2 expression cassette at the E1 locus. The ChAdOx1 adenovirus genome was excised from the BAC using unique PmeI sites flanking the adenovirus genome sequence. The virus was rescued and propagated in T-Rex 293 HEK cells (Invitrogen). Purification was by CsCl gradient ultracentrifugation. Virus titers were determined by hexon immunostaining assay and viral particles calculated based on spectrophotometry^39,40^

#### Study design animal experiments

Syrian hamsters - Syrian hamsters (4–6 weeks old, Envigo Indianapolis, IN) were vaccinated with 100 μl of 2.5 × 10^8^ infectious units of vaccine intramuscularly or 50 μl of 2.5 × 10^8^ infectious units of vaccine intranasally. Animals were vaccinated 28 days before challenge or exposure. One day prior to virus challenge or exposure animals were bled via the retro-orbital plexus. the direct challenge experiment, 10 animals per group were challenged with 40 µl of 10^4^ TCID_50_ SARS-CoV-2/human/USA/RML-7/2020 (MW127503.1) diluted in sterile Dulbecco’s Modified Eagle’s media (DMEM). In the transmission experiment, 14 unvaccinated donor animals per group were challenged with 40 µl of 10^4^ TCID_50_ SARS-CoV-2/human/USA/RML-7/2020 diluted in sterile DMEM. One day later, 14 vaccinated animals per group were co-housed with donor animals at a 2:2 or 1:1 ratio, separated by sex. Four hours later, donor animals were removed from the cage and euthanized. In each experiment, 50% of animals were male and 50% of animals were female. At 5 DPI, four animals were euthanized, and the remaining animals were followed for 21 days post challenge. Weight was recorded daily, and oropharyngeal swabs were taken daily up to 7 days post inoculation in 1 mL of DMEM supplemented with 2% fetal bovine serum, 1 mM L-glutamine, 50 U/ml penicillin and 50 μg/ml streptomycin (DMEM2). Upon euthanasia, blood and lung tissue were collected and subsequently analysed for virology and histology.

NHPs – Experimental design was based on a previously reported study^8^. Eight Indian origin rhesus macaques (5F, 3M) between 4-11 years old were sorted by sex, then by weight, and then randomly divided into two groups of four animals. Animal group size was based on initial model development^17^. The vaccine group was vaccinated with 1 ml of ChAdOx1 nCoV-19 using a MAD Nasal™ IN Mucosal Atomization Device (Teleflex, US) at −56 and −28 DPI. Within the control group, 2 animals were vaccinated via the same route with ChAdOx1 GFP, and two animals were vaccinated with ChAdOx1 GFP in 2 ml using an Omron Mesh nebulizer NE-U100. All vaccinations were done with 2.5 × 10^10^ virus particles/animal diluted in sterile PBS. Animals were challenged with SARS-CoV-2/human/USA/RML-7/2020 (MW127503.1) diluted in sterile DMEM on 0 DPI; with administration of 4 mL intratracheally and 1 mL intranasally of 2 × 10^5^ TCID_50_/mL virus suspension. Animals were scored daily by the same person who was blinded to study group allocations using a standardized scoring sheet as previously described^16^. Scoring was based on the evaluation of the following criteria: general appearance and activity, appearance of skin and coat, discharge, respiration, feces and urine output, and appetite. Clinical exams were performed on −56, −49, −42, −28, −21, −14, −7, 0, 1, 3, and 5 and 7 DPI. Nasosorption samples and blood were collected at all exam dates. Nasosorption samples were collected as previously described^41^. Briefly, a nasosorption device ((Hunt Developments UK Ltd) was inserted into the nasal cavity, and the nostril was manually held closed for 60 seconds. The swab was placed in 300 µl of AB-33K (PBS containing 1% BSA and 0.4% Tween-20) and vortexed for 30 seconds. The swab and liquid were placed on a spin filter (Agilent, 5185-5990) and spun at 16,000 rpm for 20 min. Filtered liquid was aliquoted and stored at −80°C. Nasal swabs were collected on 0, 1, 3, 5, and 7 DPI. BAL was performed on 3, 5, and 7 DPI as previously described. For each procedure, 10-30 mL of sterile saline was instilled and the sample was retrieved with manual suction.^42^ Necropsy was performed on 7 DPI and the following tissues were collected: cervical lymph node, mediastinal lymph node, nasal mucosa, trachea, all six lung lobes, right and left bronchus, spleen.

#### Cells and virus

SARS-CoV-2/human/USA/RML-7/2020 (MW127503.1) was obtained from a nasopharyngeal swab obtained on July 19, 2020. Virus propagation was performed in VeroE6 cells in DMEM2. The used virus stock was 100% identical to the initial deposited Genbank sequence and no contaminants were detected. VeroE6 cells were maintained in DMEM supplemented with 10% fetal bovine serum, 1 mM L-glutamine, 50 U/ml penicillin and 50 μg/ml streptomycin (DMEM10). VeroE6 cells were provided by Dr. Ralph Baric. Mycoplasma testing is performed at regular intervals and no mycoplasma was detected.

#### Virus titration

Tissue sections were weighed and homogenized in 750 µL of DMEM. Virus titrations were performed by end-point titration in VeroE6 cells, which were inoculated with tenfold serial dilutions of virus swab media or tissue homogenates in 96-well plates. Plates were spun down for 1 hour at 1000 rpm. When titrating tissue homogenate, cells were washed with PBS and 100 µl of DMEM2. Cells were incubated at 37°C and 5% CO2. Cytopathic effect was read 6 days later.

#### Virus neutralization

Sera were heat-inactivated (30 min, 56 °C), after which two-fold serial dilutions were prepared in DMEM2. 100 TCID_50_ of SARS-CoV-2 strain nCoV-WA1-2020 (MN985325.1) was added. After 1hr of incubation at 37°C and 5% CO_2_, the virus:serum mixture was added to VeroE6 cells. CPE was scored after 6 days at 37°C and 5% CO_2_ for 6 days. The virus neutralization titer was expressed as the reciprocal value of the highest dilution of the serum which still inhibited virus replication.

#### RNA extraction and quantitative reverse-transcription polymerase chain reaction

RNA was extracted from nasal swabs and BAL using the QiaAmp Viral RNA kit (Qiagen) according to the manufacturer’s instructions. Tissue was homogenized and extracted using the RNeasy kit (Qiagen) according to the manufacturer’s instructions. Viral gRNA^43^ and sgRNA^44^ specific assays were used for the detection of viral RNA. Five μl RNA was tested with the Rotor-GeneTM probe kit (Qiagen) or Quantstudio (Thermofisher) according to instructions of the manufacturer. Dilutions of SARS-CoV-2 standards with known genome copies were run in parallel.

#### Expression and purification of SARS-CoV-2 S and receptor binding domain

Protein production was performed as described previously^45,46^. Expression plasmids encoding the codon optimized SARS-CoV-2 full length S and RBD were obtained from Kizzmekia Corbett and Barney Graham (Vaccine Research Center, Bethesda, USA)^47^ and Florian Krammer (Icahn School of Medicine at Mt. Sinai, New York, USA)^48^. Expression was performed in Freestyle 293-F cells (Thermofisher), maintained in Freestyle 293 Expression Medium (Gibco) at 37°C and 8% CO_2_ shaking at 130 rpm. Cultures totaling 500 mL were transfected with PEI at a density of one million cells per mL. Supernatant was harvested 7 days post transfection, clarified by centrifugation and filtered through a 0.22 µM membrane. The protein was purified using Ni-NTA immobilized metal-affinity chromatography (IMAC) using Ni Sepharose 6 Fast Flow Resin (GE Lifesciences) or NiNTA Agarose (QIAGEN) and gravity flow. After elution the protein was buffer exchanged into 10 mM Tris pH8, 150 mM NaCl buffer (S) or PBS (RBD) and stored at − 80°C.

#### ELISA

ELISA was performed as described previously^8^. Briefly, maxisorp plates (Nunc) were coated overnight at 4°C with 100 ng/well S or RBD protein in PBS. Plates were blocked with 100 µl of casein in PBS (Thermo Fisher) for 1hr at RT. Serum serially diluted 2x in casein in PBS was incubated at RT for 1hr. Antibodies were detected using affinity-purified polyclonal antibody peroxidase-labeled goat-anti-monkey IgG (Seracare, 074-11-021) in casein followed by TMB 2-component peroxidase substrate (Seracare, 5120-0047). The reaction was stopped using stop solution (Seracare, 5150-0021) and read at 450 nm. All wells were washed 4x with PBST 0.1% tween in between steps. Threshold for positivity was set at 3x OD value of negative control (serum obtained from non-human primates prior to start of the experiment) or 0.2, whichever one was higher.

#### Ig subtyping and SARS-CoV-2 specific IgG/IgA quantification

Ig subtyping was performed using the isotyping Panel 1 Human/NHP Kit on the Meso Quickplex (MSD, K15203D). S and RBD antibodies were determined using the V-PLEX SARS-CoV-2 Panel 2 kit (MSD, K15383U and K15385U).

#### Antibody-dependent complement deposition

11 µl of Red FluoSpheres™ NeutrAvidin™-Labeled Microspheres (ThermoFisher, F8775) were coated with biotinylated S (25 µl at 1 mg/mL) or RBD protein (5 µl at 1 mg/mL) for 2 hours at 37°C, washed twice with PBS, and diluted in 1 mL of PBS. Serum was diluted 10x in RPMI1640 (Gibco). 10 µl of beads, 40 µl RPMI1640 and 50 ul diluted sera was mixed and incubated for 2 hours at 37°C. Guinea pig complement (Cedarlane, CL4051) was diluted 25x in gelatin veronal buffer (Boston Bioproducts, IBB-300X), 100 µl was added to the serum: bead complex and incubated at 37°C for 20 min. The serum:bead complex was then washed twice with 15 mM EDTA and incubated with 50 µl FITC-conjugated-anti-C3 antibody (100x in PBS, MP Biomedical, 855385) for 15 min at RT in dark. Serum:bead complexes were washed three times with PBS and analyzed on a BD FACS Symphony A5 (BD Biosciences) flow cytometer using a high throughput sampler within 1 hour of completion of protocol. All samples were run in duplicate. Serum:bead complexes were gated by FSC vs SSC to remove debris, followed by red bead fluorescence gating in the PE channel, and then the geometric mean fluorescent intensity (MFI) in the FITC channel was determined using FlowJo 10 (BD Biosciences) software and analyzed in Graphpad Prism version 8.3.0..

#### Antibody dependent monocyte cellular phagocytosis

Beads were prepared as described above. Serum was diluted 100x in RPMI1640, 100 µl was mixed with 10 µl beads and incubated at 37°C for 2 hours. Beads were washed once with RPMI1640. THP-1 cells (ATCC, TIB-202) were diluted to 1.25 × 10^5^ cells/mL in RPMI1640, 100 µl was added per sample, and incubated at 37°C for 18 hours. Cells were fixed in 10% formalin for 15 min at RT in dark, washed twice with PBS and ran on a BD FACS Symphony A5 (BD Biosciences) flow cytometer using a high throughput sampler. All samples were run in duplicate as described above.

#### Antibody-dependent NK cell activation

NK-cell activation was assessed using methods similar to those previously described.1,2 Briefly, cells were isolated from 30 mL of heparin-treated whole blood collected from a healthy human donor (NIH IRB 01-I-N055) using the RosetteSepTM Human NK Cell Enrichment Cocktail according to the manufacturer’s instructions (Stem Cell). NK cells were rested overnight at 37°C in complete RPMI 1640 media supplemented with 10% fetal bovine serum and 1 ng/mL of IL-15 (Stem Cell). Nunc MaxiSorpTM 96-well ELISA plates (Thermo Fisher) were coated with 3 µg/mL of SARS-CoV-2 S or RBD antigen for 2 hours at 37°C. Plates were subsequently washed and blocked with a solution of 5% BSA in 1X DPBS overnight at 4oC.

Sera samples collected at −56 and 0 DPI were diluted 1:25 in blocking buffer, plated in duplicates, and incubated on the coated ELISA plates for 2 hours at 37°C. NK cells were resuspended in a staining cocktail containing anti-CD107a-PE/Cy7 antibody (BioLegend), GolgiStop (BD), and GolgiPlug (BD). After removal of sera from the plate, 5.0×10^4^ NK cells were added per well and incubated at 37°C for 6 hours.

Surface staining was carried out using anti-CD56-BUV737 (BD), anti-CD16-BV510 (BioLegend), and anti-CD3-BV650 (BD) antibodies prior to fixation and permeabilization using Cytofix/CytoPermTM solution (BD). Intracellular staining was performed using anti-IFNγ-PerCP/Cy5.5 (BioLegend) and anti-MIP-1β-PE (BD) antibodies. Data acquisition was performed using FACSymphonyTM A5 (BD). NK cells were identified by gating on CD3-CD16+ CD56+ cells.

#### Integrated analysis of multivariate antibody and virology profiles

Principal component analysis was performed using the R packages “FactoMineR” and “factoextra” to compare antibody and virology profiles. Spearman rank (two-sided) correlation coefficients for pairwise comparisons between all variables were generated using the R “cor” function; the correlation matrix was visualized in R using “ggcorplot.”

#### cDNA Synthesis

cDNAs were prepared according to Briese *et al*.^49^ Briefly, RNA was extracted from hamster swabs and tissues following the QiaAmp Viral RNA extraction protocol (Qiagen, Germantown, MD) and 11 µL was taken into the SuperScript IV First-Strand cDNA synthesis system (ThermoFisher Scientific, Waltham, MA) following the manufacture’s recommendations. After RNase H treatment, second-strand synthesis was performed using Klenow fragment (New England Biolabs, Ipswich, MA) following the manufacturer’s recommendations. The resulting double-stranded cDNAs (ds-cDNA) were then purified using Ampure XP bead purification (Beckman Coulter, Pasadena, CA) and eluted into 30 µL water.

#### Sequencing Library Construction and SARS-CoV2 Enrichment

To construct sequencing libraries, 25 µL ds-cDNA was brought to a final volume of 53 µL in Elution Buffer (Agilent Technologies, Santa Clara, CA) and sheared on the Covaris LE220 (Covaris, Woburn, MA) to generate an average size of 180-220 bp. The following settings were used: peak incident power, 450 watts; duty factor, 15%; cycles per burst, 1000; and time, 300 seconds. The Kapa HyperPrep kit was utilized to prepare libraries from 50 µL of each sheared cDNA sample following modifications of the Kapa HyperPrep kit, version 8.20, and SeqCap EZ HyperCap Workflow, version 2.3, user guides (Roche Sequencing Solutions, Inc., Pleasanton, CA). Adapter ligation was performed for 1 hour at 200C using the Kapa Unique-Dual Indexed Adapters diluted to 1.5 µM concentration (Roche Sequencing Solutions, Inc., Pleasanton, CA). Following ligation, samples were purified with AmPure XP beads (Beckman Coulter, Brea, CA) and subjected to double-sided size selection as specified in the SeqCap EZ HyperCap Workflow User’s guide. Pre-capture PCR amplification was performed using 12 cycles, followed by purification using AmPure XP beads.

Purified libraries were assessed for quality on the Bioanalyzer 2100 using the High Sensitivity DNA chip assay (Agilent Technologies, Santa Clara, CA). Quantification of pre-capture libraries was performed using the Qubit dsDNA HS Assay kit and the Qubit 3.0 fluorometer following the manufacturer’s instructions (ThermoFisher Scientific, Waltham, MA).

The myBaits Expert Virus bait library was used to enrich samples for SARS-CoV-2 according to the myBaits Hybridization Capture for Targeted NGS, version 4.01, protocol. Briefly, libraries were sorted according to estimated genome copies and pooled to create a combined mass of 2 µg for each capture reaction. Depending on estimated genome copies, two to six libraries were pooled for each capture reaction. Capture hybridizations were performed 16-19 hours at 650C and subjected to 8-14 PCR cycles after enrichment. SARS-CoV-2-enriched libraries were purified and quantified using the Kapa Library Quant Universal qPCR mix in accordance with the manufacturer’s instructions. Libraries were diluted to a final working concentration of 1-2 nM, titrated to 20 pM, and sequenced as 2 − 150 bp reads on the MiSeq sequencing instrument using the MiSeq Micro kit version 2 (Illumina, San Diego, CA).

#### Next Generation Sequencing data analysis

Raw fastq reads were adapter trimmed using Cutadapt v 1.12^50^, followed by quality trimming and quality filtering using the FASTX Toolkit (Hannon Lab, CSHL). Reads were paired up and aligned to the SARS-CoV-2 genome from isolate SARS-CoV-2/human/USA/RML-7/2020 (MW127503.1) using Bowtie2 v 2.2.9^51^. PCR duplicates were removed using Picard MarkDuplicates v 2.18.7 (Broad Institute). Variant detection was performed using GATK HaplotypeCaller v 4.1.2.0^52^ with ploidy set to 2. Raw variant calls were filtered for high confidence variants using bcftools filter^53^ with parameters QUAL > 500 and DP > 20.

#### Histology and immunohistochemistry

Necropsies and tissue sampling were performed according to IBC-approved protocols. Lungs were perfused with 10% neutral-buffered formalin and fixed for eight days. Hereafter, tissue was embedded in paraffin, processed using a VIP-6 Tissue Tek (Sakura Finetek, USA) tissue processor, and embedded in Ultraffin paraffin polymer (Cancer Diagnostics, Durham, NC). Samples were sectioned at 5 µm, and resulting slides were stained with hematoxylin and eosin. an in-house SARS-CoV-2 nucleocapsid protein rabbit antibody (Genscript) at a 1:1000 dilution was utilized to detect specific anti-CoV immunoreactivity, carried out on a Discovery ULTRA automated staining instrument (Roche Tissue Diagnostics) with a Discovery ChromoMap DAB (Ventana Medical Systems) kit. The tissue slides were examined by a board-certified veterinary anatomic pathologist blinded to study group allocations. 18 sections, taken from six different lung lobes are evaluated for each animal; a representative lesion from each group was selected for the figure.

#### Statistics

Two-tailed Mann–Whitney tests, two-way ANOVA, mixed-effect analysis, Fisher test, Spearman rank (two-sided) correlation coefficients, or Kruskall-Wallis analysis were conducted to compare differences between groups using Graphpad Prism version 8.3.0. Statistical tests used are identified in figure legends or main text.

**Table S1.** Pathology and IHC scoring direct challenge hamsters. H&E was scored as follows: 0 = not present; 1 = 1-10%; 2 = 11-25%; 3 = 26-50%; 4 = 51-75%; 5 = 76-100%. IHC was scored as follows: 0 = not present; 1 = rare/few; 2 = scattered; 3 = moderate; 4 = numerous; 5 = diffuse.

|  |  | IN-vaccinated animals |  |  |  | IM-vaccinated animals |  |  |  | Control animals |  |  |  |
| --- | --- | --- | --- | --- | --- | --- | --- | --- | --- | --- | --- | --- | --- |
| H&E | Lesions % | 0 | 0 | 0 | 0 | 0 | 0 | 0 | 0 | 70 | 60 | 40 | 70 |
|  | Interstitial pneumonia | 0 | 0 | 0 | 0 | 0 | 0 | 0 | 0 | 4 | 4 | 3 | 4 |
|  | Bronchiolitis | 0 | 0 | 0 | 0 | 0 | 0 | 0 | 0 | 2 | 2 | 3 | 3 |
|  | Alveolar exudate | 0 | 0 | 0 | 0 | 0 | 0 | 0 | 0 | 2 | 2 | 2 | 2 |
|  | Type II pneumocyte hyperplasia | 0 | 0 | 0 | 0 | 0 | 0 | 0 | 0 | 2 | 2 | 3 | 2 |
|  | Perivascular leukocyte infiltration | 0 | 0 | 0 | 0 | 0 | 0 | 0 | 0 | 2 | 2 | 2 | 2 |
|  | Edema | 0 | 0 | 0 | 0 | 0 | 0 | 0 | 0 | 2 | 2 | 1 | 3 |
| IHC | Staining % | 0 | 0 | 0 | 0 | 0 | 0 | 0 | 0 | 70 | 60 | 20 | 70 |
|  | Type I and II pneumocytes | 0 | 0 | 0 | 0 | 0 | 0 | 0 | 0 | 4 | 4 | 3 | 4 |
|  | Exudate | 0 | 0 | 0 | 0 | 0 | 0 | 0 | 0 | 1 | 0 | 1 | 1 |
|  | Bronchiolar epithelium | 0 | 0 | 0 | 0 | 0 | 0 | 0 | 0 | 3 | 1 | 2 | 3 |

**Table S2.** Pathology and IHC scoring transmission hamsters. H&E was scored as follows: 0 = not present; 1 = 1-10%; 2 = 11-25%; 3 = 26-50%; 4 = 51-75%; 5 = 76-100%. IHC was scored as follows: 0 = not present; 1 = rare/few; 2 = scattered; 3 = moderate; 4 = numerous; 5 = diffuse.

|  |  | IN-vaccinated animals |  |  |  | IM-vaccinated animals |  |  |  | Control animals |  |  |  |
| --- | --- | --- | --- | --- | --- | --- | --- | --- | --- | --- | --- | --- | --- |
| H&E | Lesions % | 0 | 0 | 0 | 0 | 0 | 20 | 10 | 5 | 50 | 50 | 50 | 40 |
|  | Interstitial pneumonia | 0 | 0 | 0 | 0 | 0 | 3 | 3 | 2 | 3 | 3 | 3 | 3 |
|  | Bronchiolitis | 0 | 0 | 0 | 0 | 0 | 0 | 0 | 0 | 3 | 2 | 2 | 2 |
|  | Alveolar exudate | 0 | 0 | 0 | 0 | 0 | 2 | 1 | 2 | 2 | 3 | 2 | 2 |
|  | Type II pneumocyte hyperplasia | 0 | 0 | 0 | 0 | 0 | 3 | 1 | 0 | 2 | 2 | 1 | 1 |
|  | Perivascular leukocyte infiltration | 0 | 0 | 0 | 0 | 0 | 0 | 2 | 0 | 1 | 1 | 1 | 2 |
|  | Edema | 0 | 0 | 0 | 0 | 0 | 0 | 0 | 0 | 3 | 1 | 2 | 2 |
| IHC | Staining % | 0 | 0 | 0 | 0 | 0 | 5 | 5 | 5 | 60 | 60 | 50 | 30 |
|  | Type I and II pneumocytes | 0 | 0 | 0 | 0 | 0 | 1 | 1 | 1 | 4 | 4 | 4 | 2 |
|  | Exudate | 0 | 0 | 0 | 0 | 0 | 2 | 0 | 0 | 1 | 1 | 1 | 1 |
|  | Bronchiolar epithelium | 0 | 0 | 0 | 0 | 0 | 1 | 1 | 0 | 3 | 1 | 2 | 3 |

**Figure S1.**
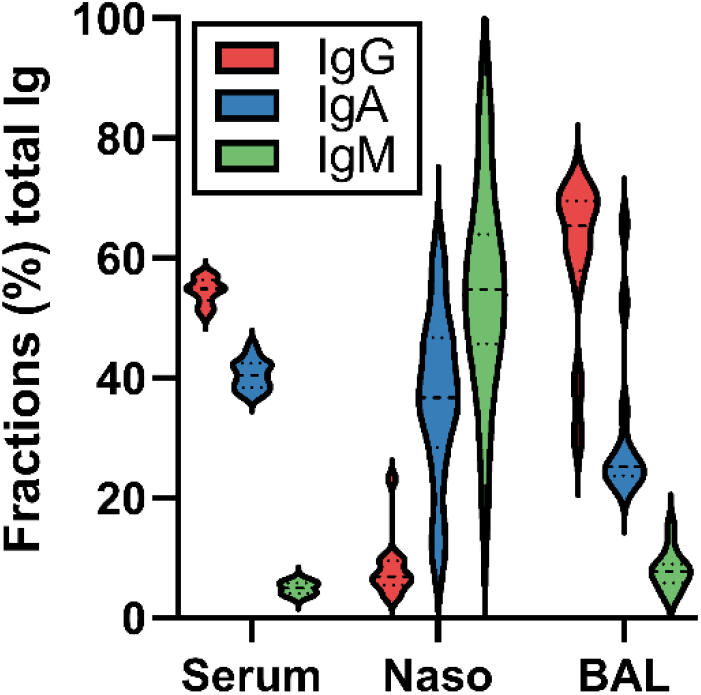
Fractions of IgA, IgG, and IgM in serum, nasosorption or BAL samples. Twenty-four serum, 14 nasosorption, and 12 BAL samples were investigated.

**Figure S2.**
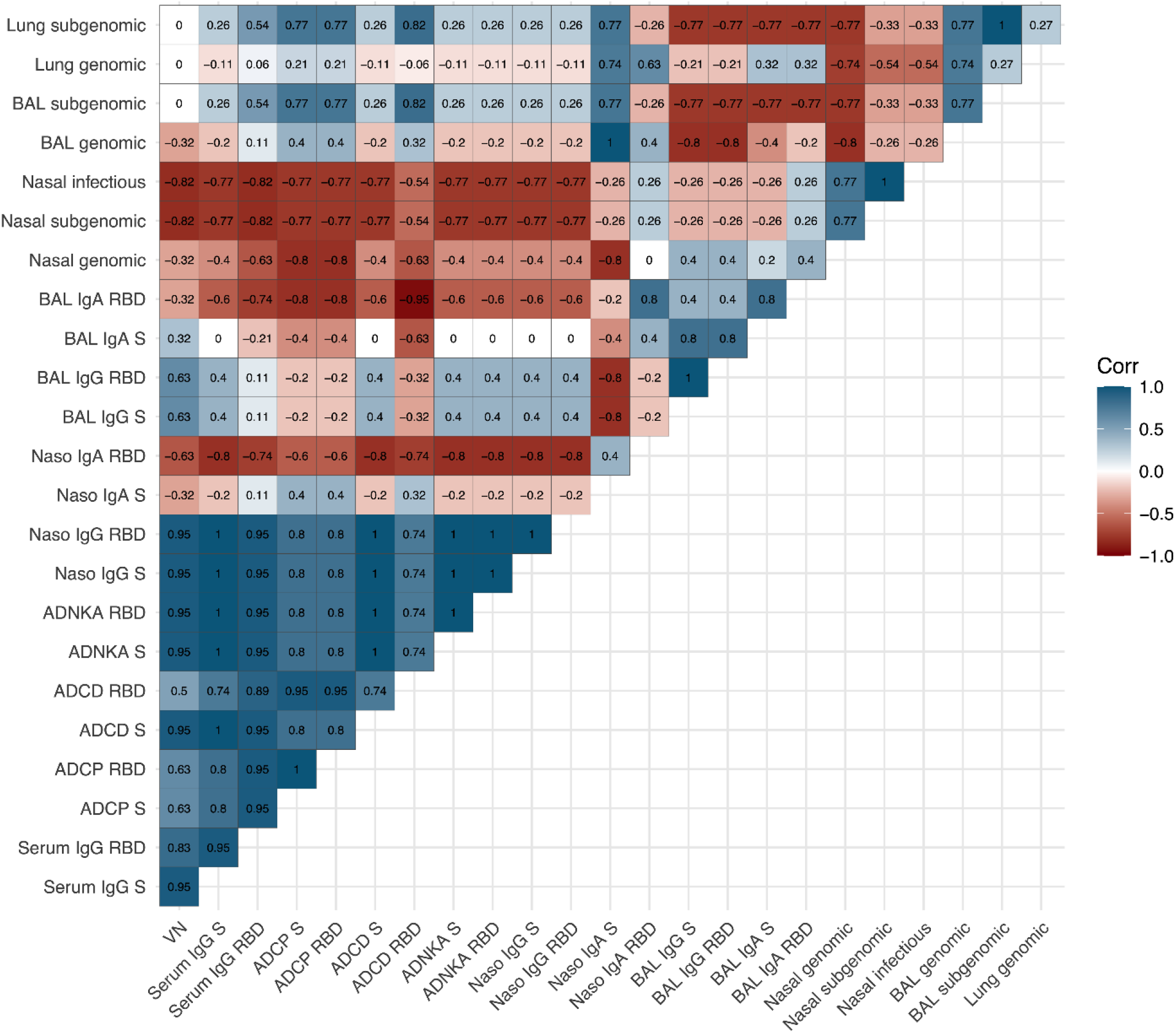
Correlation matrix featuring all immunology and virology measures. Correlation heatmap, depicted as a matrix, representing pairwise correlations between all antibody and virology variables in IN-vaccinated animals. The two-sided Spearman rank correlation coefficient is indicated within each square.

